# Phylogenetic and spatial determinants of leaf endophyte microbiomes in the flowering plant genus *Heuchera* (Saxifragaceae)

**DOI:** 10.1101/2023.05.23.541975

**Authors:** Dexcem J. Pantinople, Reagan Conner, Stephanie Sutton-Dauber, Kelli Broussard, Carolina M. Siniscalchi, Nicholas J. Engle-Wrye, Heather R. Jordan, Ryan A. Folk

**Affiliations:** Department of Biological Sciences, 295 Lee Boulevard, Mississippi State, MS 39762, USA

**Keywords:** bacteria, community assembly, fungi, *Heuchera*, leaf endophyte, microbial diversity, plant microbiome, plant-microbe interactions, Saxifragaceae

## Abstract

*Premise*: Endophytic plant-microbe interactions range from mutualistic relationships that confer important ecological and agricultural traits to neutral or quasi-parasitic relationships. In contrast to root-associated endophytes, the role of environmental and host-related factors for acquiring leaf endophyte communities remains relatively unexplored. Here we assess leaf endophyte diversity to test the hypothesis that membership of these microbial communities is driven primarily by abiotic environment and host phylogeny.

*Methods:* We used a broad geographic coverage of North America in the genus, *Heuchera* (Saxifragaceae). Bacterial and fungal communities were characterized with 16S and ITS amplicon sequencing, using QIIME2 to call operational taxonomic units and calculate species richness, Shannon diversity, and phylogenetic diversity. We assembled environmental predictors for microbial diversity at collection sites including latitude, elevation, temperature, precipitation, and soil parameters.

*Results:* We find differing assembly patterns for bacterial and fungal endophytes; we found that only host phylogeny is significantly associated with bacteria, while geographic distance alone was the best predictor of fungal community composition. Species richness and phylogenetic diversity are very similar across sites and species, with only fungi showing a response to aridity and precipitation for some metrics. Unlike what has been observed with root-associated microbial communities, in this system microbes show no relationship with pH or other soil factors.

*Conclusions:* Host phylogeny and geographic distance independently influence different microbial communities, while aridity and precipitation determine fungal diversity within leaves of *Heuchera*. Our results indicate the importance of detailed clade-based investigation of microbiomes and the complexity of microbiome assembly within specific plant organs.

## INTRODUCTION

Endophytic plant-microbe interactions are common to all land plants, which are host to a diverse range of microbial assemblages, including bacteria, archaea, fungi, and protists. Endophytes, microorganisms that spend all or a portion of their lifetime within plant tissues (Hardoim et al., 2015), confer such positive functional capacity as abiotic stress response, growth promotion, life history traits, and pathogen or herbivore defense, as well as the potential for negative interactions approaching pathogenic relationships (Hardoim et al., 2008; Khare et al., 2018; Dini-Andreote, 2020; Trivedi et al., 2020; O’Brien et al., 2021). A context-dependent switch between positive and negative interactions in many endophytic systems means plant endophytes form an excellent system for understanding the formation and maintenance of mutualisms (Eaton et al., 2011). In addition, multiple variables including, host and endophyte physicochemical characteristics, biotic and abiotic factors, and the microbial dynamics within the microbiome affect the nature of these associations (Hardoim et al., 2015).

Endophytic relationships are relatively well-characterized in several economically important species such as major pasture grasses (Clay, 1990; Leuchtmann, 1992; Schardl and Tsai, 1992) and crop plants (Fisher and Petrini, 1992; Fisher et al., 1992; Larran et al., 2002; Comby et al., 2016; Correa-Galeote et al., 2018), mostly investigated under regulated experimental conditions. In natural environments, endophyte diversity surveys have been conducted primarily at broad phylogenetic (Yeoh et al., 2017) and geographic scales (Yang et al., 2019). These natural surveys, primarily focused on root-associated microbiomes, show broadly that soil properties are the most important drivers of plant-associated microbiome diversity, much as in free-living soil microbiomes (Thompson et al., 2017; Bahram et al., 2018). Nevertheless, host plant phylogeny plays an important and incompletely characterized subsidiary role for both bacterial and fungal communities, a role possibly rooted in shared evolutionary history or conserved plant host traits (Yeoh et al., 2017; Yang et al., 2019). An evolutionary host effect on endophytes may indicate either (1) functional selection of associated microbes by the plant (or vice versa) or (2) shared coevolutionary history between plants and their endophytes. Since we know that global diversity patterns show strong mismatches between plants and free-living microbes (Cameron et al., 2019), there also exists the strong potential for conflict between drivers of distribution and diversity between endophytes and their hosts when plant-microbe associations are particularly intimate.

By contrast to root endophytes and rhizosphere associates, the role of potential external and host-driven factors for assembling leaf endophyte communities remains relatively unexplored. The leaf ecosystem still lags behind other tissue types in endophyte research despite supporting a wide variety of microbial communities and having a total surface area that is roughly twice that of Earth (Vorholt, 2012; Harrison and Griffin, 2020). This leads to the prediction that leaf endophyte communities should be more insulated from the soil environment because of the more controlled environment of internal leaf tissues across varying soil substrates, especially in contrast to rhizosphere communities. Composition of foliar endophyte communities should then have a limited response to soil ecology but a stronger response to climatic and other similar abiotic factors. Moreover, aboveground conditions that leaves encounter are unlikely to affect soil environments (Monteith and Unsworth, 2008). However, a strong case exists for potential host phylogenetic constraints on leaf endophyte communities due to phylogenetically conserved differences in leaf tissue traits across taxa (Tellez et al., 2022) as well as the potential for vertical transmission (particularly well-characterized in grasses; Schardl, 2001; Bright and Bulgheresi, 2010) and semi-vertical transmission with hosts through primarily within-population sources of infection (Frank et al., 2017; Kandel et al., 2017).

A study system that can link across population-level and phylogenetic scales (Graham et al., 2018) would provide insight into how plant-microbe interactions arise and particularly insight into the phylogenetic level at which host specificity is relevant. Such a multi-scale view would also link phylogenetically broad and single-species surveys performed to date. As advocated by (Jung et al., 2021), multi-scale research is also important for generating genotype × environment viewpoints on plant microbiomes and giving researchers additional power to dissect factors that promote different microbiome assemblages and result in gradients in plant-microbe interactions.

Here, we take a novel approach that uses broad geographic coverage of North America within the restricted phylogenetic scope of a recent radiation. Using the host system *Heuchera*, a cliff-dwelling genus of flowering plants in the family Saxifragaceae with well-characterized phylogenetic relationships and habitat specialization patterns across the genus (Folk et al., 2017; Folk, Visger, et al., 2018), we leverage strong phylogenetic and population sampling to explicitly assess diversity trends at multiple evolutionary levels, from phylogenetic to within-population diversity. We assembled a series of predictors via global environmental layers, including elevation, temperature, precipitation, soil parameters, and latitude. We use multiple assessments of leaf endophyte diversity to (1) test the hypothesis that these communities, in contrast to root-associated microbiome, are defined primarily by non-edaphic abiotic environmental variables, and (2) by host phylogeny. Finally (3), we assess both bacterial and fungal endophyte components to ask whether these communities are shaped by distinct environmental factors.

## MATERIALS AND METHODS

### Host organism

*Heuchera* is a genus of approximately 45 species of flowering plants in Saxifragaceae that is endemic to rock outcrops and montane areas in North America. It occurs from sea level to ∼4000 m of elevation across broad temperate environmental gradients including temperate deciduous and evergreen woodland, plains, high alpine scree, and chaparral. Edaphic variation is also high and ranges from strong calciphile taxa (e.g., *H. longiflora*) to some of the most acidic substrates in North America (*H. parviflora* var. *saurensis*), with many narrow endemics particular to specific rock substrates. Hence, this genus forms a robust system for evaluating plant-microbe interactions across the strong, continent-level environmental gradients. Aside from small numbers of taxa included in broad surveys (e.g., Jumpponen and Trappe, 1998; Zhang and Yao, 2015) and characterizations of arbuscular mycorrhizae (Anneberg and Segraves, 2019), endophytic microbial associates are currently unknown for the family Saxifragaceae.

### Sampling

We began with broad species-level sampling across the study group, including 40 out of 64 currently recognized specific and subspecific taxa (65%). Taxa covered are geographically representative of the range of the genus north of Mexico (Fig. 1) and include all recognized sections (Folk, 2015). In addition to this broad phylogenetic-aware sampling of the host plant genus, we leveraged population-level sampling from two previous studies on host plant phylogeography in the *Heuchera parviflora* species complex (Folk and Freudenstein, 2015) and the *H. longiflora* complex (Folk et al., 2018), as well as new sampling performed for this study in the *H. americana* × *H. richardsonii* hybrid zone (see Wells, 1984). The newly sampled taxa were: *H. americana* group: *H. americana* var. *americana*, *H. americana* var. *hirsuticaulis*, *H. richardsonii*; *H. longiflora* group: *H. longiflora* var. *aceroides*, *H. longiflora* var. *longiflora*; *H. parviflora* group: *H. missouriensis, H. parviflora* var. *parviflora, H. parviflora* var. *saurensis, H. puberula*. Sampling is summarized in Fig. 1 and Appendix S1 (See Supplemental Data with this article).

**Fig. 1.**
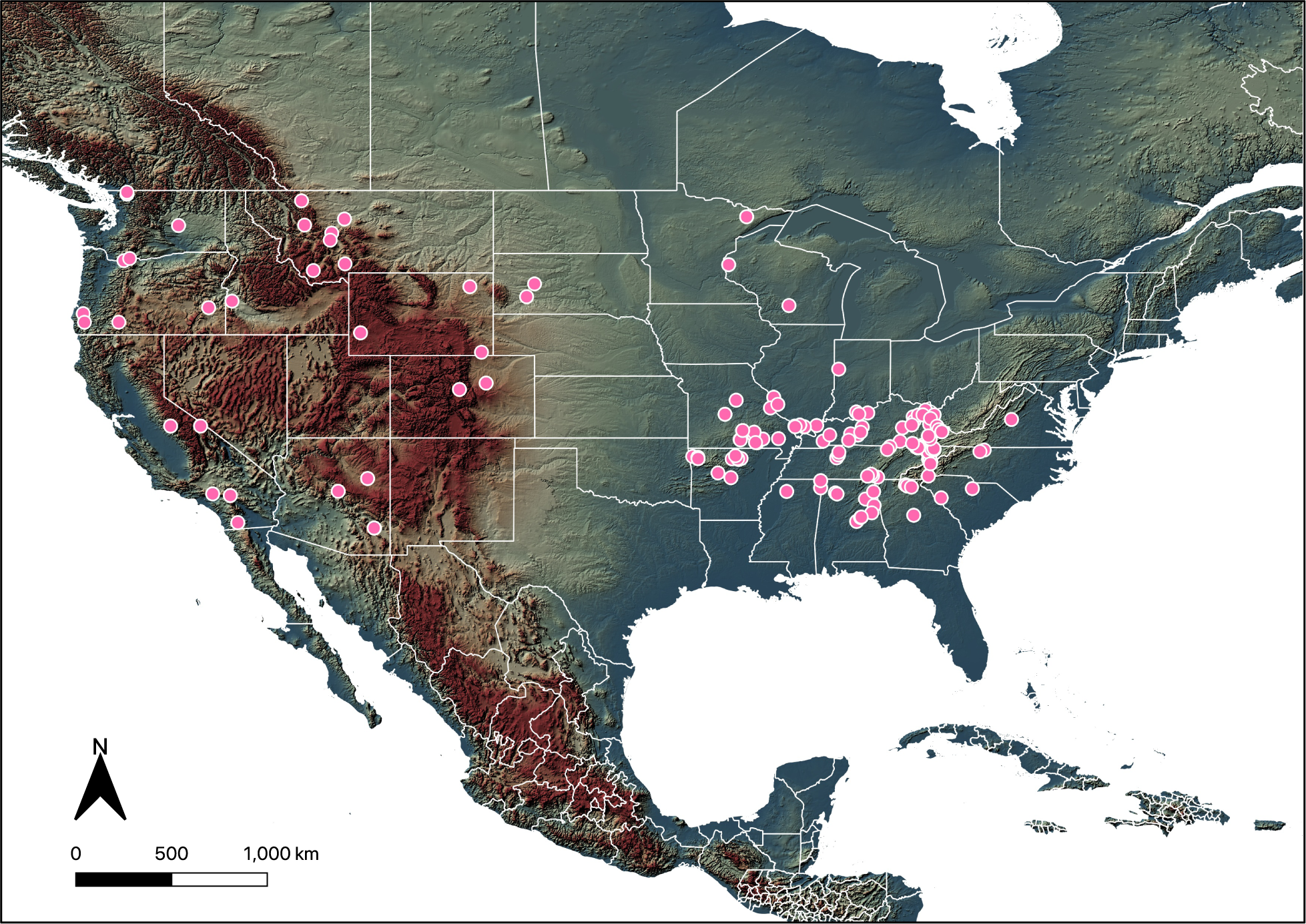
Map of *Heuchera* samples used in the study (pink circles). Map generated using the QGIS Software (v3.24; QGIS Development Team 2021).

### DNA extraction

Plant materials were either rapidly frozen at -80°C and subsequently dehydrated or primarily dried in silica-gel prior to extraction. For DNA extraction, we chose 20-30 mg of tissue without visible lesions or other obvious disease symptoms. The tissue was incubated for 1 min each in 70% molecular-grade ethanol and 5% bleach to disrupt and eliminate DNA of potential epiphytic microbes, respectively. Tissues were then washed twice in molecular-grade water to remove residual bleach and homogenized with metal beads in a Fisherbrand Bead Mill 24 homogenizer (Fisher Scientific, Waltham, Massachusetts, USA). We extracted DNA with a standard CTAB protocol (Doyle and Doyle, 1987) with the addition of 90 mg ascorbic acid and 100 mg polyvinylpyrrolidone-40 (PVP-40) per extraction to eliminate plant secondary compounds, per previous optimizations on this plant material (Folk and Freudenstein, 2014). Finally, all extractions were cleaned using a silica column (GeneJET PCR purification kit, ThermoScientific, Waltham, Massachusetts, USA) per manufacturer instructions and extractions were quantified with a Qubit 4 fluorometer using Qubit Broad Range assay reagents.

### Amplification methods

We used two different amplicon sequencing approaches to characterize both bacterial and fungal communities. Bacterial sequencing was validated in-house using primers 515f and 806r from the Earth Microbiome Project (Thompson et al., 2017) targeting the V4 region of 16S ribosomal DNA and the following thermocycler protocol: initial denaturation at 95°C for 3 mins, then 35 cycles of: (95°C for 45 s, annealing at 52°C for 1 min, and 72°C for 1.5 mins), then a final elongation step of 72°C for 10 mins. Successful amplicons were concentration-normalized and sent to the Michigan State RSTF core for sequencing 250 bp paired-end reads on an Illumina MiSeq using a one-step amplification protocol (Kozich et al., 2013). All amplification steps used DreamTAQ Mastermix (Thermo Fisher, Waltham, Massachusetts, USA), primer concentrations of 0.5 µM except as noted below, and were performed with filter pipette tips under a dedicated PCR hood that was bleach- and UV-sterilized before each use to minimize contamination.

Fungal characterization used the ITS1 region and the primers ITS1FI2 and ITS2 from (Schmidt et al., 2013). To verify the presence of amplifiable DNA, we first validated the presence of the desired product using the primers directly and the following thermocycler protocol: initial denaturation at 95°C for 3 mins, then 35 cycles of: (95°C for 45 s, annealing at 50°C for 1 min, and 72°C for 1 min), then a final elongation step of 72°C for 10 mins. We then re-amplified successful samples from total DNA using ITS1FI2 and ITS2 primers that were tagged with 5’ end overhangs specified by the sequencing center using the following thermocycler protocol: initial denaturation at 95°C for 5 mins, then 30 cycles of: (95°C for 30 s, annealing at 52°C for 30 s, and 72°C for 30 s), then a final elongation step of 72°C for 5 mins; primers for this reaction were at 0.1 µM. Successful amplicons were submitted to the Michigan State RSTF core for a second barcoding amplification and sequencing. Sequencing instrumentation and wet lab precautions followed those for 16S (above).

### Sequence processing

We performed sequence analyses within the QIIME 2 package (Caporaso et al., 2010; Bolyen et al., 2019). Reads were first denoised via Dada2 (Callahan et al., 2016) in order to error-correct and merge paired-end reads and remove sequence chimeras. As part of this step, primers were trimmed from the 5’ end and, based on Phred quality plots in FastQC (Andrews, 2015), 50 bp were trimmed from the 3’ end of the R2 reads.

For taxonomic classification, we used the Greengenes database (McDonald et al., 2012) for bacterial 16S reads, and the UNITE database (Nilsson et al., 2019) for fungal ITS reads, following recommendations in the QIIME documentation for preparing the taxonomic classifier via a naive Bayesian approach (QIIME module fit-classifier-naive-bayes). We clustered the Greengenes database at 97% and UNITE at 99% identity. We then performed taxonomic classifications of the merged reads against these databases using QIIME module sklearn (Pedregosa et al., 2011). For endophyte tissues, 16S and ITS amplicon sequencing approaches were expected to generate host plant DNA sequences due to off-target amplification of organellar 16S rDNA and nuclear ITS, respectively. Based on extensive optimizations, we implemented separate strategies for efficiently removing host DNA from each of these genetic loci. For 16S, we removed host DNA using annotated chloroplast and mitochondrial OTU classifications from the Greengenes taxonomy (level 3 [class] and level 5 [family], respectively). For ITS, we customized the UNITE database by adding host plant ITS sequences we have previously generated (Folk and Freudenstein, 2014), and removed host sequences based on level 6 (genus) OTU classifications.

### Environmental predictor assembly

We used global interpolated datasets to infer environmental factors at each collection locality. The variables used and sources were: Mean Annual Temperature (measured in °C) and Annual Precipitation (mm; BIOCLIM, (Hijmans et al., 2005)), aridity (see below), elevation (m; GTOPO30, https://www.usgs.gov/centers/eros/science/usgs-eros-archive-digital-elevation-global-30-arc-second-elevation-gtopo30), soil pH, sand percent, and carbon content (the last measured in permilles; SoilGrids, Hengl et al., 2017). An aridity index was calculated as precipitation / potential evapotranspiration (see Middleton et al., 1992) using data from WorldClim2 and Envirem (Fick and Hijmans, 2017; Title and Bemmels, 2018). Note that this aridity index *decreases* with increasing aridity; arid conditions are generally those with index values < 0.5. Environmental values were associated with geolocated sampling localities using scripts published previously (https://github.com/ryanafolk/Saxifragales_spatial_scripts/tree/master/Extract_point_values). Finally, given that varying latitudinal gradients in diversity have been documented for soil (Bahram et al., 2018) and marine microbes (Ibarbalz et al., 2019), we also directly used the latitude of our collecting localities as a predictor.

### Community diversity

We used QIIME to generate two primary descriptors of community diversity. First, we characterized measures of overall diversity using Shannon Entropy, a diversity measure that includes both taxon presence-absence information and abundance. We then calculated Faith’s PD, which represents the sum of phylogenetic branch lengths connecting a microbial community. We applied these diversity metrics to only the three species groups with strong population sampling to enable comparisons among host taxa with replicate sampling. Given the presence of high levels of host DNA despite a high sequencing effort in some samples (Results) and relatively low endophyte diversity per sample (Results and also see Bulgarelli et al., 2013), sequence rarefaction was set to 11 to include as many samples as possible.

We used both a generalized linear model (GLM) and a linear mixed-modeling (LMM) framework (R package lmer) to understand how these diversity statistics relate separately to environmental drivers and host identity. All environmental predictors, as well as latitude, were included as fixed model terms in both model classes. Host plant species taxonomy was also included as a random term in the LMM to separately partition variation attributable to host taxon. We used the step function (R package lmerTest) to perform model selection via AIC and calculate predictor significance using an automated backwards approach. The AIC model selection favored GLM as the optimal fit model given our observed data. Analyses were performed using R Statistical Software (v4.1.2; R Core Team 2021).

### Community composition

In order to characterize differences among communities in terms of taxon composition, we used the UniFrac distance metric, which accounts both for shared taxon presence/absence and for phylogenetic branch length, here including all samples. We used a Mantel testing approach to ask whether matrices of UniFrac distance were associated with each of either geographic distance, environment, or host phylogenetic distance.

Environment distances were Euclidean distances on the environmental predictors, where two matrices were prepared segregating the environmental predictors into soil and non-soil factors. Since geographic and environmental distances were strongly correlated, we additionally used a partial Mantel approach to control environmental factors for geography. Host phylogenetic distances were patristic distances calculated from the host plant phylogeny of Folk et al. (2017); this was a phylogenetic estimate based on phylogenomic data with complete species-level sampling of the host plants used here. Since that previous phylogeny did not include population-level sampling, population samples were imputed by placing them within the phylogeny based on taxonomic identifications and assuming zero within-taxon branch lengths. Analyses were performed using R Statistical Software (v4.1.2; R Core Team 2021).

## RESULTS

### Sequencing

For 16S sequencing, we recovered a mean of 236,938 reads per sample across 139 successful samples, with 1,737 total bacterial OTUs across all samples and a mean of 97% host DNA prevalence. The 5 most dominant bacterial phyla by decreasing order of prevalence were Proteobacteria (6 to 100% per sample), Bacteroidetes (0 to 83%), Actinobacteria (0 to 38%), Verrucomicrobia (0 to 13%), and Cyanobacteria (0 to 48%) (Fig. 2A). Finer level classifications of OTUs recovered largely corresponded to typical endophytes documented elsewhere, such as, in decreasing order of overall prevalence for 16S: *Sphingomonas* (which reached highest prevalence at up to 100%), Comamonadaceae, Chitinophagaceae, *Methylobacterium*, *Blastomonas, Hymenobacter, Pseudomonas,* and Opitutaceae. Similar to other surveys in natural populations (Yeoh et al., 2017), potential diazotrophs (genera *Rhizobium, Bradyrhizobium, Mesorhizobium, Frankia*) were observed at low frequencies (up to 8% of total 16S reads) in almost all samples (Appendix S2).

**Fig. 2.**
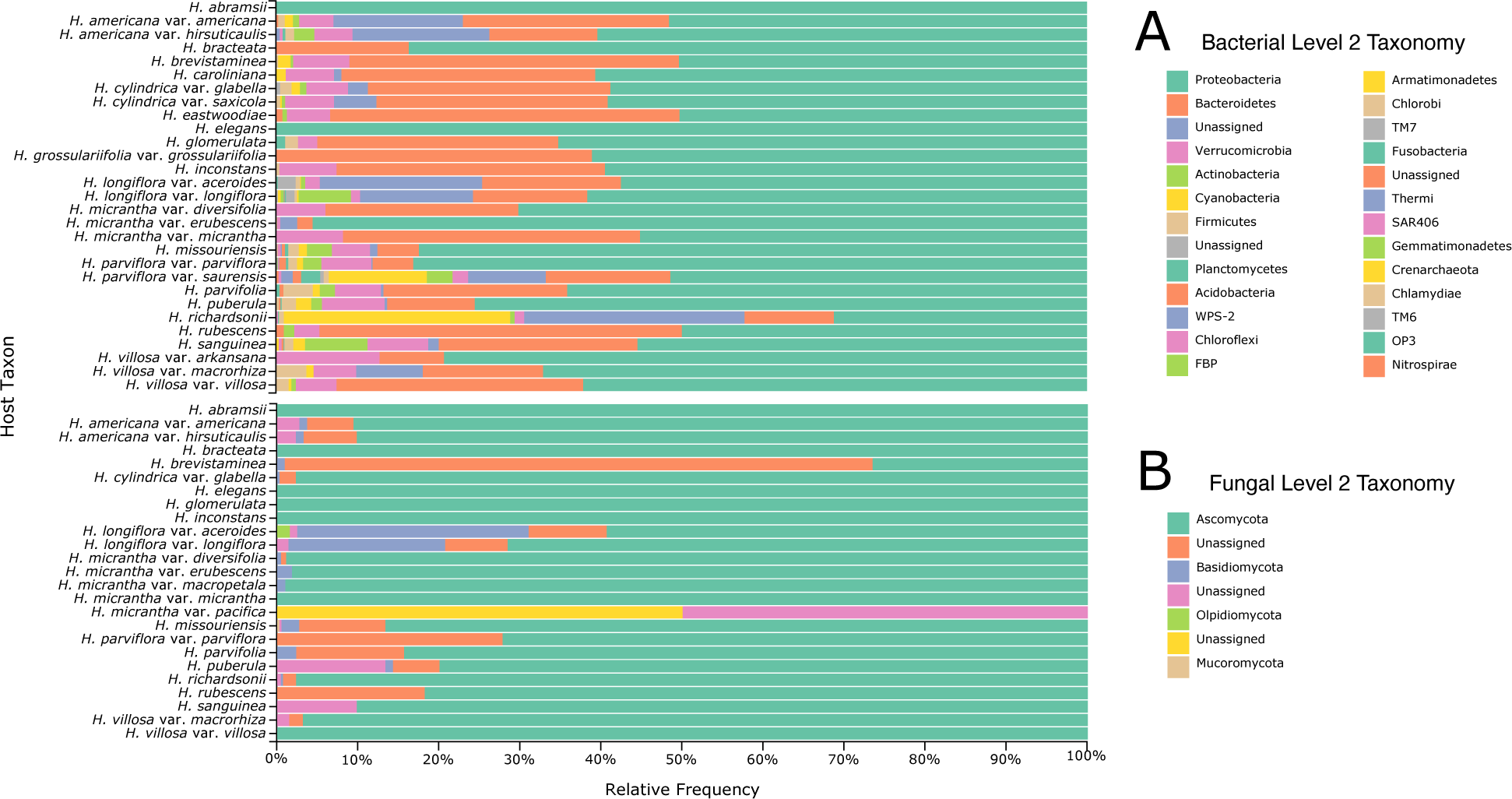
(A) Bacterial and (B) fungal endophyte phylum-level diversity and relative abundance per

For ITS sequencing, we recovered a mean of 185,997 reads per sample across 133 successful samples, with a total of 1,082 fungal OTUs and a mean of 99% host DNA prevalence; lower fungal diversity compared to bacterial diversity has been previously documented in leaf endophytes (Bulgarelli et al., 2013). By far the most dominant phylum was Ascomycota (only missing in a single sample; otherwise, 5 to 100%), with Basidiomycota (0–44%, absent in a slight majority of samples), Olpidiomycota (0 to 77%, absent in most samples), and Mucoromycota (<1%) as minor community members. (Fig. 2B) As with bacteria, fungal fine-level OTU designations generally contain previously documented endophytes; in order of decreasing abundance the most prevalent were *Penicillium,* Pleosporaceae*, Septoria,* and *Alternaria* (all four up to 100% abundance)*, Mycosphaerella, Tetracladium, Ramularia,* and *Colletotrichum* (Appendix S3).

### Leaf endophyte diversity patterns

Using a mixed-model framework, we tested for a role of climate, soil environment, latitude, elevation, and species identity on leaf endophyte diversity as measured by Shannon entropy and Faith’s phylogenetic diversity. For bacteria, we found the null model was favored for both diversity metrics, meaning leaf endophyte diversity metrics were insensitive to the predictors we measured. However, for fungi, aridity and precipitation were significant drivers of Shannon diversity for fungal endophytes (*P* = 0.001676, 0.003246 respectively), while the null model was favored for Faith’s PD (although aridity index was marginally significant; *P* = 0.0508; Table 1). Based on examination of boxplots (Fig. 3), the only species group that had a clear trend in Shannon diversity or Faith’s PD was the *H. parviflora* group, although this difference was not significant (16S: ANOVA, *F*_3,26_ = 1.604, 1.797, *P* = 0.212, 0.173 respectively; ITS: *F*_2,19_ = 1.528, 1.071, *P* = 0.242, 0.362 respectively; Fig. 3); taxa in the other two species groups had near-identical means.

**Fig. 3.**
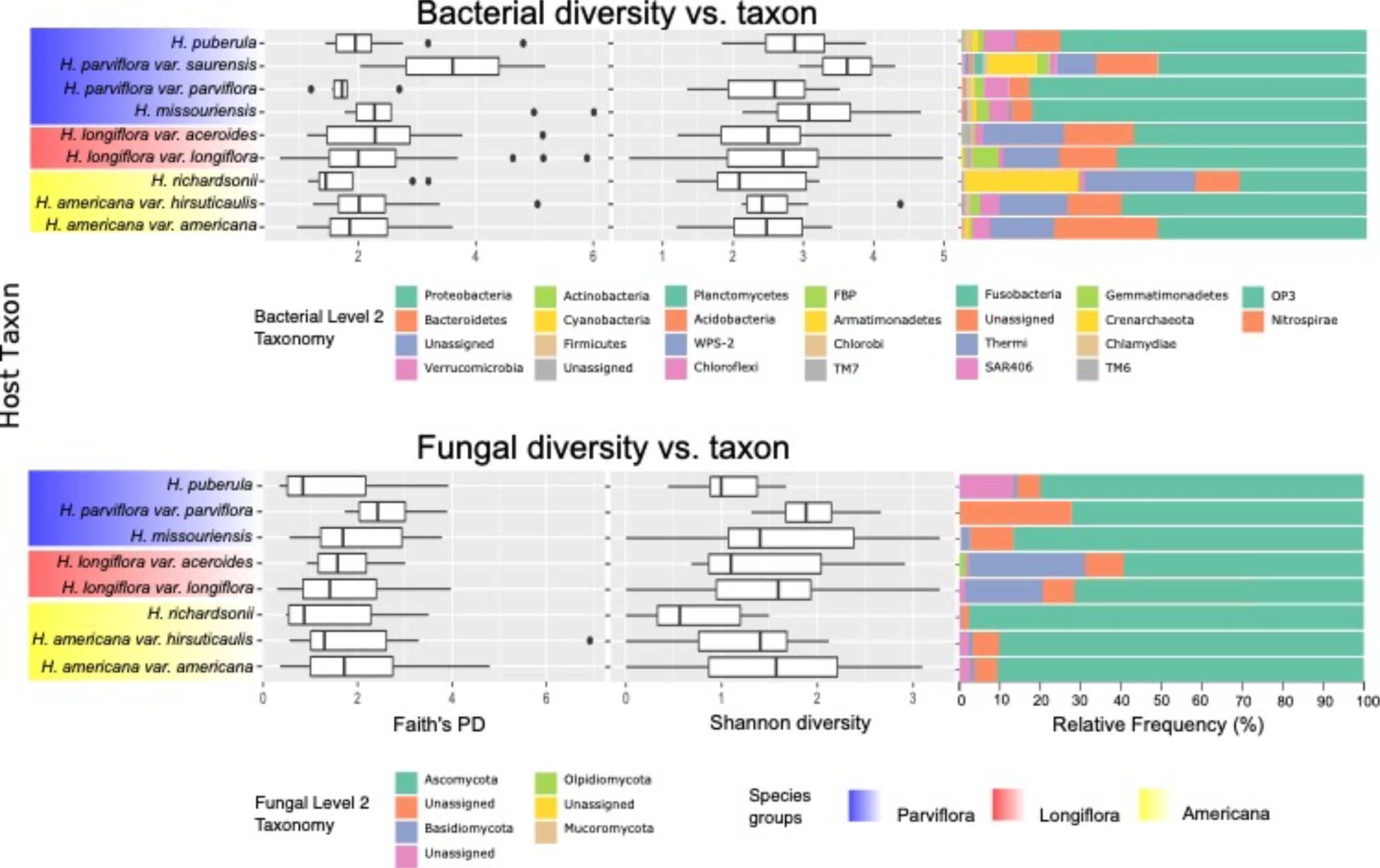
Boxplot of microbial endophyte Faith’s Phylogenetic and Shannon Diversity with relative abundance across strongly sampled host taxa.

**Table 1.**
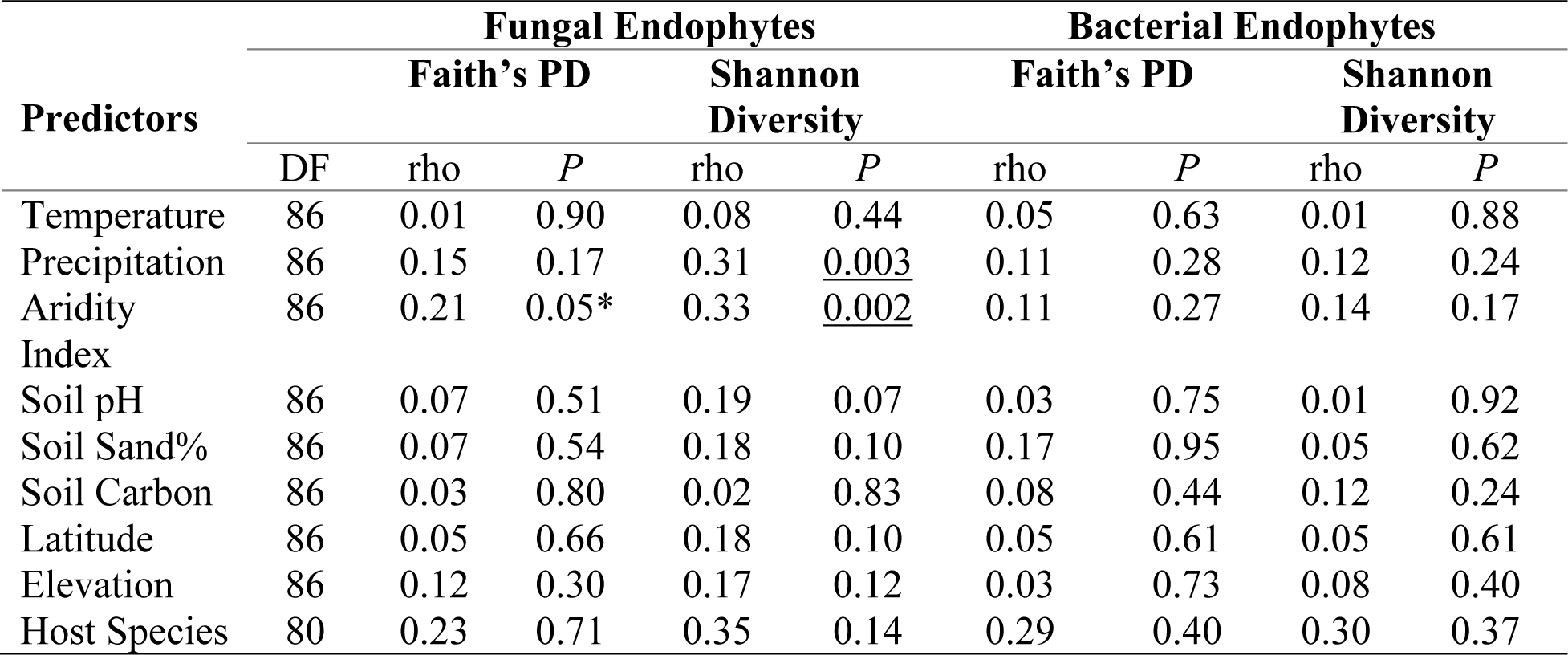
Correlation between predictors and leaf endophyte diversity metrics; significant values (*P* < 0.05) are underlined; *marginally significant.

| Predictors | Fungal Endophytes |  |  |  |  | Bacterial Endophytes |  |  |  |
| --- | --- | --- | --- | --- | --- | --- | --- | --- | --- |
|  | Faith's PD |  |  | Shannon Diversity |  | Faith's PD |  | Shannon Diversity |  |
|  | DF | rho | *P* | rho | *P* | rho | *P* | rho | *P* |
| Temperature | 86 | 0.01 | 0.90 | 0.08 | 0.44 | 0.05 | 0.63 | 0.01 | 0.88 |
| Precipitation | 86 | 0.15 | 0.17 | 0.31 | <u>0.003</u> | 0.11 | 0.28 | 0.12 | 0.24 |
| Aridity Index | 86 | 0.21 | 0.05* | 0.33 | <u>0.002</u> | 0.11 | 0.27 | 0.14 | 0.17 |
| Soil pH | 86 | 0.07 | 0.51 | 0.19 | 0.07 | 0.03 | 0.75 | 0.01 | 0.92 |
| Soil Sand% | 86 | 0.07 | 0.54 | 0.18 | 0.10 | 0.17 | 0.95 | 0.05 | 0.62 |
| Soil Carbon | 86 | 0.03 | 0.80 | 0.02 | 0.83 | 0.08 | 0.44 | 0.12 | 0.24 |
| Latitude | 86 | 0.05 | 0.66 | 0.18 | 0.10 | 0.05 | 0.61 | 0.05 | 0.61 |
| Elevation | 86 | 0.12 | 0.30 | 0.17 | 0.12 | 0.03 | 0.73 | 0.08 | 0.40 |
| Host Species | 80 | 0.23 | 0.71 | 0.35 | 0.14 | 0.29 | 0.40 | 0.30 | 0.37 |

### Leaf endophyte community composition

Using UniFrac distances as a characterization of leaf endophyte community composition, we asked whether communities were associated with any of three potential drivers: geography (that is, isolation-by-distance), soil or non-soil environment, or host phylogeny. For bacteria, we found that only host phylogeny was significant (Mantel test, *P* = 0.00229977; Table 2). For fungi, we found that both geography (Mantel test, *P*<0.001) and both soil and non-soil environment (Mantel test, *P* = 0.00209979, 0.00849915) were significantly associated with UniFrac distance. Given that we found spatial autocorrelation among both sets of environmental predictors (Mantel test, both *P* < 0.001), we controlled for geography using a partial Mantel approach. We found after this correction that soil was marginally significant (*P* = 0.047695) while non-soil environment was no longer significant (*P* = 0.26197) for fungi, indicating that geography was the best predictor of fungal diversity and the effect of environment independent of geography was weak.

**Table 2.** Microbial endophyte partial Mantel tests *P*-values; significant values (*P*<0.05) are underlined; *marginally significant.

|  | Fungal Endophytes | Bacterial Endophytes |
| --- | --- | --- |
| Geographic distance | <u>0.0005</u> | 0.56 |
| Host phylogeny | 0.62 | <u>0.002</u> |
| Soil environment | 0.05* | 0.12 |
| Climate | 0.26 | 0.72 |

## DISCUSSION

Our investigation of the leaf endophyte microbiome in *Heuchera* provides a first report on the phylogenetic and environmental determinants associated with leaf microbiome diversity and community assembly at a broad phylogenetic and geographic scale using culture-independent approaches. The foliar microbial endophytes we recovered from *Heuchera* generally matched those found in other leaf endophyte studies. Proteobacteria, Bacteroidetes, and Actinobacteria have consistently been reported as dominant and prevalent members of endophytic bacterial assemblages within plant tissues (Hardoim et al., 2015; Zarraonaindia et al., 2015; Coleman-Derr et al., 2016; de Souza et al., 2016; Ding and Melcher, 2016; Aydogan et al., 2018; Wemheuer et al., 2019; Mina et al., 2020; Yang et al., 2023). At the genus level, *Pseudomonas*, *Sphingomonas*, *Methylobacterium*, and *Hymenobacter* were also found to be relatively abundant in leaves of both cultivated (Hallmann et al., 1997; Rosenblueth and Martínez-Romero, 2006; Miliute et al., 2015; Afzal et al., 2019; Christian et al., 2021) and non-cultivated plants (Ding and Melcher, 2016; Afzal et al., 2019). On the other hand, the majority of leaf endophytic fungi in *Heuchera* belonged to Ascomycota and Basidiomycota, similarly reported as two of the most dominant fungal endophyte classes in its close relative, *Saxifraga* (Zhang and Yao, 2015) and across a variety of host plants (Zimmerman and Vitousek, 2012; Jin et al., 2013; Fan et al., 2020; Pang et al., 2022). In addition, *Penicillium*, Pleosporaceae, *Alternaria*, and *Colletotrichum* have also been documented as predominant fungal endophytes associated with leaves of multiple host plant species (Fisher et al., 1992; Araújo et al., 2001; Gamboa and Bayman, 2001; Romero et al., 2001; Douanla-Meli et al., 2013; Jin et al., 2013; Matsumura and Fukuda, 2013; Fang et al., 2019).

### Environment and endophyte diversity

Assessing microbial diversity patterns, we found that bacterial endophyte species (Shannon’s index) and phylogenetic (Faith’s PD) diversity were remarkably consistent across host species and all environmental variables measured. Fungal endophyte diversity, however, was significantly greater in less arid and high precipitation regions (although insignificant in multivariate analyses), which is in accordance with observations of increased richness of foliar endophytic fungi of an annual grass at wetter locations in the Mediterranean by Penner and Sapir (2021), as well as of a tree species in a Hawaiian terrain by Zimmerman and Vitousek (2012). Our results showing non-significance of latitude contrast with previous investigations demonstrating a commonly observed latitudinal diversity gradient, in which diversity declines from equatorial to polar regions. For example, Arnold and Lutzoni (2007) showed that diversity of foliar fungal endophytes follows the classical pattern of increasing diversity towards tropical areas (Canada to Panama). On the other hand, there is almost no knowledge regarding bacterial endophyte diversity patterns across latitudinal gradients. Our study therefore represents a primary demonstration of a non-significant pattern of foliar bacterial endophyte species and phylogenetic diversity across a relatively broad latitudinal range. Moreover, climate, elevation, and soil environment were weak predictors of the diversity of foliar bacterial endophytes in *Heuchera.* This pattern is consistent with previous works across host plants in which abiotic factors have little to no influence on leaf bacterial richness and composition. For example, several studies have shown that precipitation generally does not exert a significant effect on bacterial diversity (Hirano et al., 1996; Copeland et al., 2015; Stone and Jackson, 2019, 2021; Wemheuer et al., 2020). Wemheuer and colleagues (2020) also reported no significant correlation of bacterial endophyte diversity with temperature and elevation in *Theobroma cacao* leaves (also true with fungi, except temperature).

Thus, microbial leaf endophyte diversity in *Heuchera* is generally robust to differences in the abiotic environment. There may be several non-exclusive reasons for this. First, the internal leaf tissue may provide a more stable environment, insulating the effects of constant changes occurring in the surrounding environment. Second, differences of our observations from the results of previous works may be attributed to the broader phylogenetic and geographic scale of our research, extensive host species and population sampling in natural environments. Lastly, taxa we studied may also influence results, as different host taxa may have differing microbial interactions across varying environmental conditions.

### Effect of host phylogeny

Our investigation on the factors associated with leaf endophyte recruitment revealed that host phylogeny alone significantly influences bacterial community structure, while fungal composition was best predicted by geographical location. Several more focused studies have reported similar patterns, demonstrating that leaf endophytic bacterial communities are chiefly controlled by host identity (Ding et al., 2013; Mina et al., 2020), as well as showing that host biogeography and other abiotic factors play a minor role in bacterial community assembly (Coleman-Derr et al., 2016). Fungal endophyte communities, on the other hand, have been suggested to show similar patterns as our observations. For example, foliar fungal endophyte community structure was found to be strongly correlated with geographic distance in several oak species, showing similarities of fungal communities between species from adjacent sites, regardless of host habitat and phylogeny, as well as changes in climatic and environmental conditions (Collado et al., 1999; Lau et al., 2013; Koide et al., 2017). Biogeography was also a primary influence on foliar fungal endophyte community recruitment across several plant hosts including species of *Agave* (Coleman-Derr et al., 2016), and conifers (Langenfeld et al., 2013).

### Geographic distance

Our observation that isolation-by-distance was significant for fungi and not bacteria is a remarkable parallel to recent global-scale work on soil microbiomes (Bahram et al., 2018), where both environmental parameters and geographic distance significantly determined fungal diversity. This contrasting pattern has also been previously revealed by multiple comparative investigations, reporting distinct drivers of microbiome community composition between bacteria and fungi, specifically with fungal community assembly being influenced by geographic distance more than bacterial communities (Shakya et al., 2013; Coleman-Derr et al., 2016; Wei et al., 2022). This similarity in findings across disparate plant organs and taxa may reflect distinct dispersal ecologies of fungi and bacteria. Foliar fungal endophytes are usually horizontally transmitted as spores or small pieces of hyphae via air (Rodriguez et al., 2009), which suggests that geographic location plays a significant role in endophytic community recruitment. Dispersal limitation may be one of the possible explanations for this phenomenon. For instance, Zhang and colleagues (2021) found strong evidence supporting the ‘size-dispersal’ hypothesis demonstrating that larger fungi are more dispersal constrained than smaller bacterial microorganisms. This can lead to geographic heterogeneity of fungal endophyte communities and as a result, community similarity declines with growing geographic distance. Our results for bacteria, on the other hand, suggest a level of host control over bacterial community colonization of internal plant tissues. This may be attributed to varying internal physical, physiological, and biochemical environment across species of *Heuchera*, as well as specific host plant genotype traits that act as habitat filters to select for distinct microbial community species.

### Edaphic ecology

We also demonstrate here that the *Heuchera* leaf endophyte microbiome shows no relationship with the soil environment, a contrast to what has been observed in rhizosphere and root endophyte communities (Fierer and Jackson, 2006; Baker et al., 2009; Afzal et al., 2011; Bokati et al., 2016). Van Bael and colleagues (2017) similarly suggest that soil environment gradients do not significantly influence foliar endophyte diversity and community assembly. This may be due to the buffering of edaphic conditions in the more insulated internal leaf environment of the host where microbial communities inhabit. Indeed, in a recent work by Zhou et al. (2023), soil salinity determined endophytic bacterial communities in roots but not in leaves, where host leaf metabolism has more control over community assembly.

It is, however, important to note that observed patterns in this study may not hold true across the plant kingdom or to even broader geographic ranges. Multiple studies have shown contrasting patterns (e.g., Gomes et al., 2018; Wemheuer et al., 2019; Shen et al., 2020; Brigham et al., 2023), suggesting that leaf bacterial and fungal endophyte community structure are probably driven by multiple different factors including geographic location, host characteristic, soil environment, climatic and other abiotic and biotic variables. In addition, the influence of these factors may be especially dependent on the taxa being investigated, the geographic and sampling scale of the study, and the locality.

### DNA sourcing

Our work also derives substantially from silica-dried collections, an approach used previously to characterize legume nodule microbiomes (Johnson, 2019). That we recover as major community components numerous bacteria and fungi genera previously known to be typical plant endophytes indicates that useful insights can be derived from diverse preservation strategies. Easy-to-use preservation approaches are especially suitable for widely spread and inaccessible field sites for broad geographic surveys. Herbarium materials prepared under less controlled conditions than those used here have been the subject of several studies; (Daru et al., 2018; Bieker et al., 2020) were able to obtain useful endophyte microbiome data from herbarium specimens, although with higher quantities of exogenous DNA due to inconsistent mounting and storage procedures. However, materials from herbaria may prove useful in future studies to track how endophytic communities might change through time. In addition to herbaria, large, preserved tissue resources exist in several museums and other institutions as well as individual labs that would, together with a similar approach to ecological predictor assembly via georeferences, enable broad-scale surveys of endophyte diversity potentially beyond the scale of purpose-collected microbial materials.

## CONCLUSION

The significance of environmental and host-related factors in driving the assembly of leaf endophyte communities has received comparatively less attention in comparison to more extensive research on root and rhizosphere endophytes. Here, we applied a broad geographic and phylogenetic sampling to assess leaf endophyte diversity, testing the hypothesis that these communities are primarily driven by host phylogeny and abiotic environment. Our results revealed differing community assembly patterns for bacterial and fungal endophytes. We found that only host phylogeny significantly influences bacterial endophyte composition, while geographic distance was the most important determinant of endophytic fungal communities. Moreover, endophyte diversity patterns were found to be consistent across sites and host species, with only fungal diversity being significantly greater in less arid and high precipitation regions for some metrics. The present study also introduces silica-dried collection as an effective and efficient preservation approach for broad-scale leaf microbiome studies. Our findings highlight the value of in-depth clade-based microbiome research and the intricacy of microbiome assembly within certain plant organs.

## ACKNOWLEDGEMENTS

We thank the SRI Fund (College of Arts and Sciences, Mississippi State University) for financial support in optimizing wet lab procedures.

## AUTHOR CONTRIBUTIONS

RAF conceived this study with the assistance of DJP; NJE-W and RAF conducted the fieldwork; RC, SS-D, and KB performed the wet lab work, in consultation with CMS and HRJ; DJP and RAF analyzed the data and wrote the manuscript. All authors contributed to all drafts and gave final approval for publication.

## DATA AVAILABILITY STATEMENT

All sequence data and raw reads are deposited in the Sequencing Read Archive, under BioProject XXX. Scripts used for implementing QIIME, diversity, and statistical analyses are available on GitHub (https://github.com/dexcemp/heuchera_microbiome).

## ORCID

Dexcem Pantinople https://orcid.org/0009-0003-7522-464X

Carolina Siniscalchi https://orcid.org/0000-0003-3349-5081

Nicholas Engle-Wrye https://orcid.org/0000-0003-1374-0895

Heather Jordan https://orcid.org/0000-0002-4197-2194

Ryan Folk https://orcid.org/0000-0002-5333-9273

## SUPPORTING INFORMATION

Additional supporting information may be found online in the Supporting Information section at the end of the article.

**Appendix S1.**
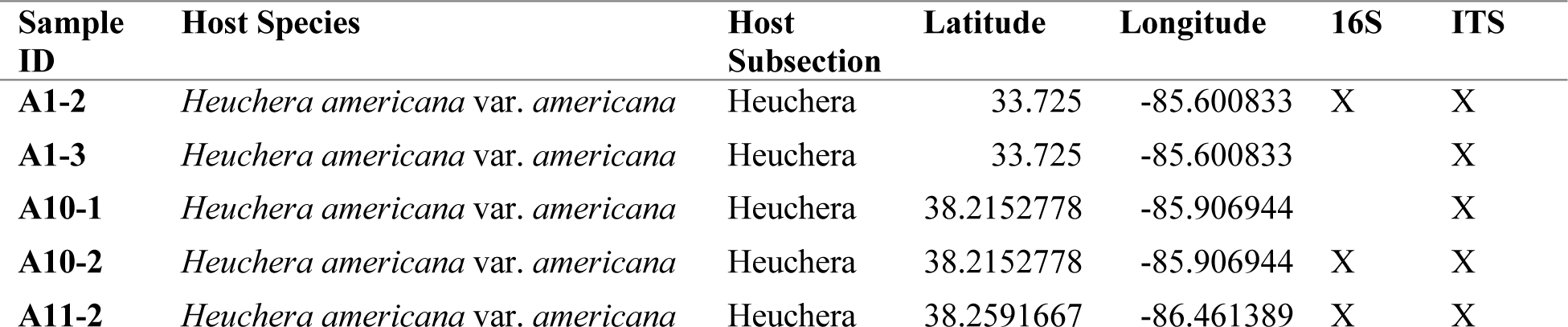
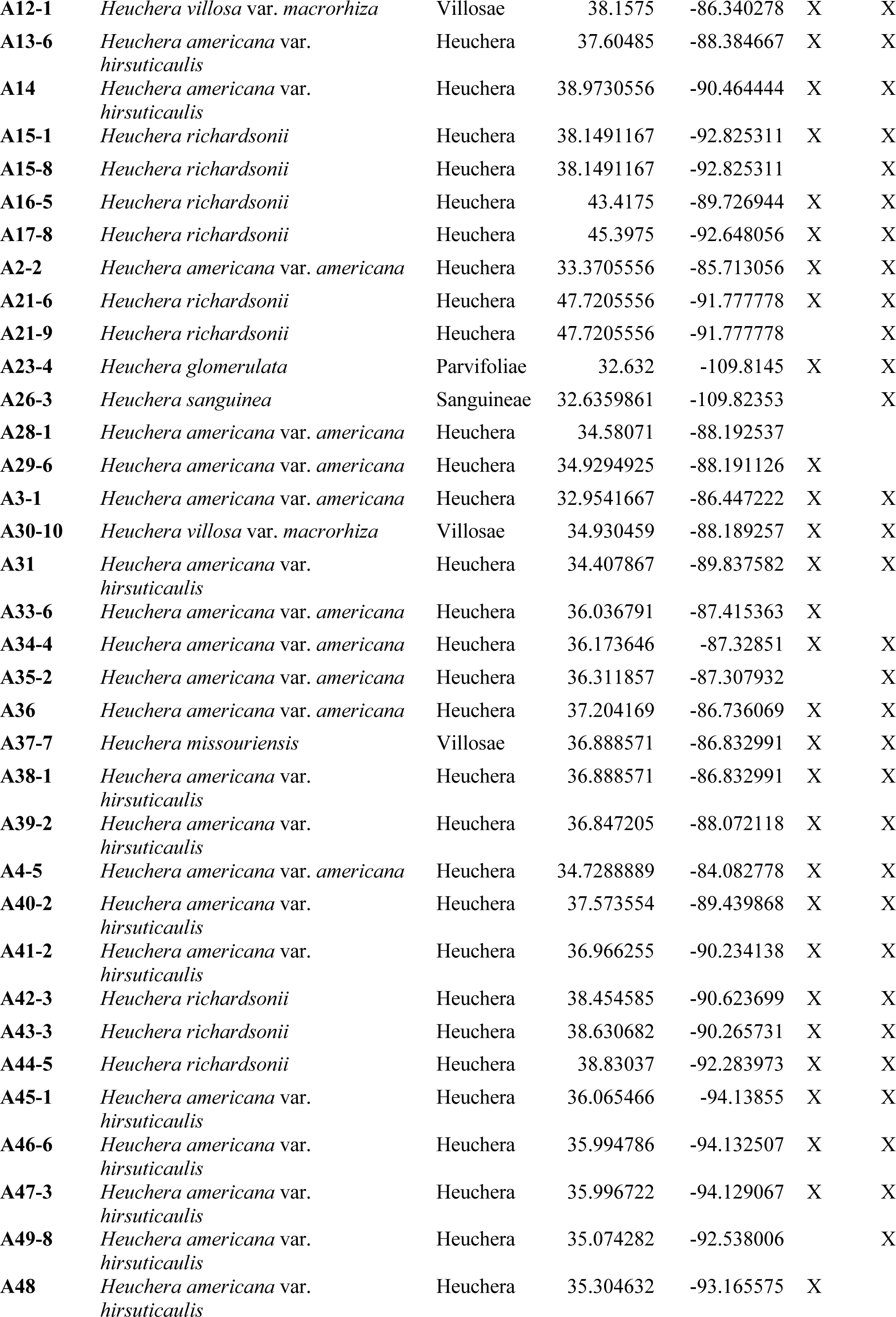
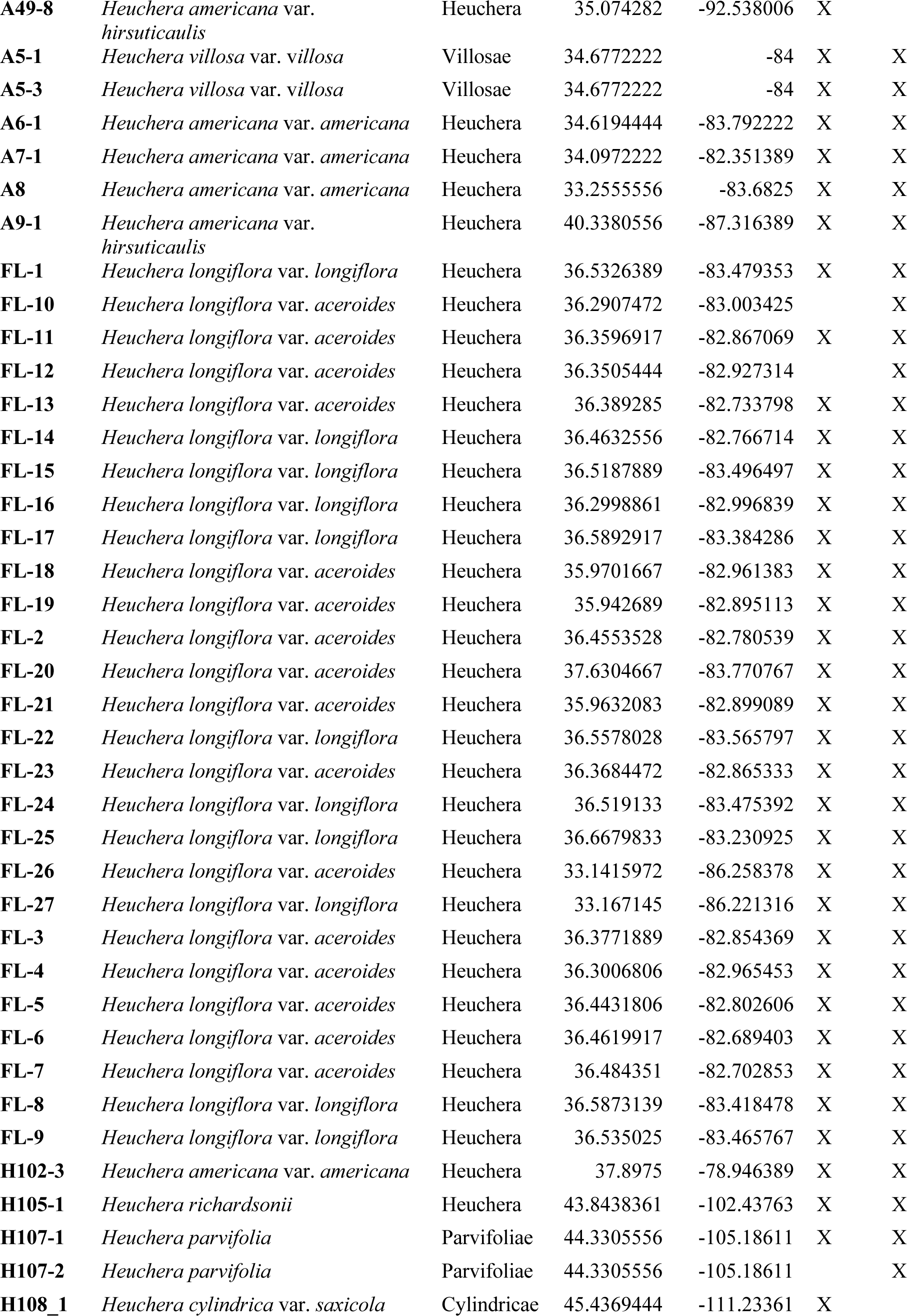
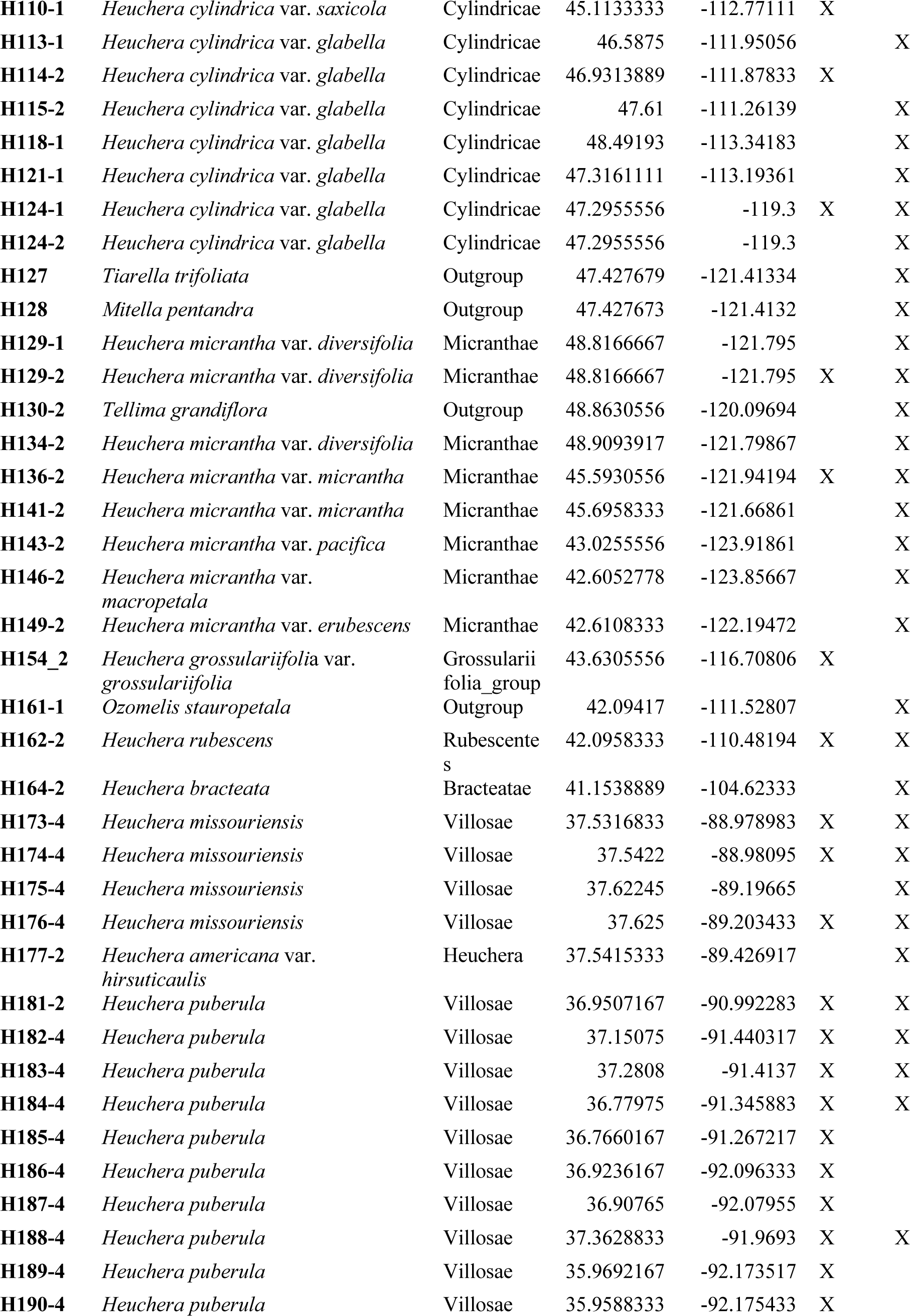
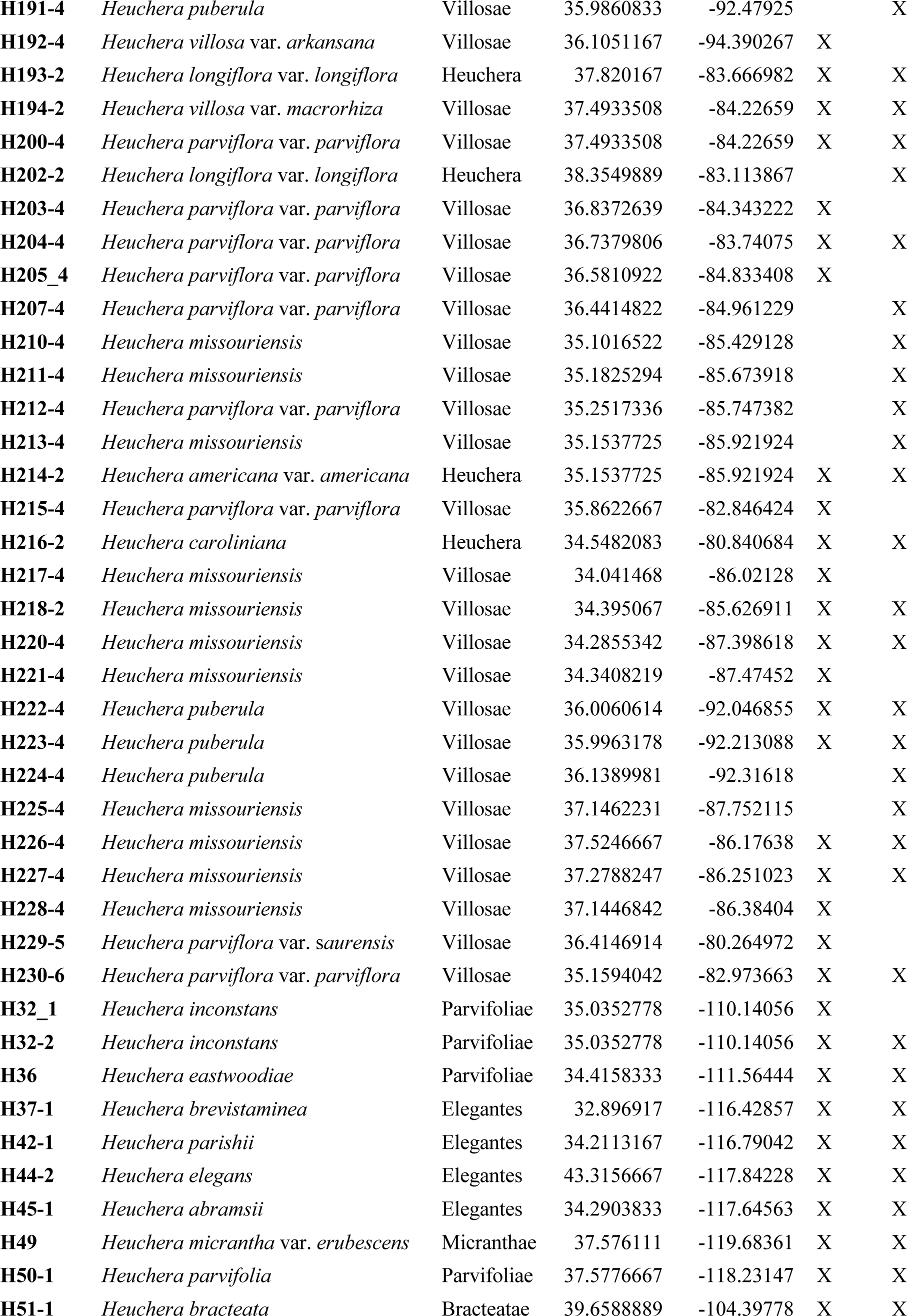
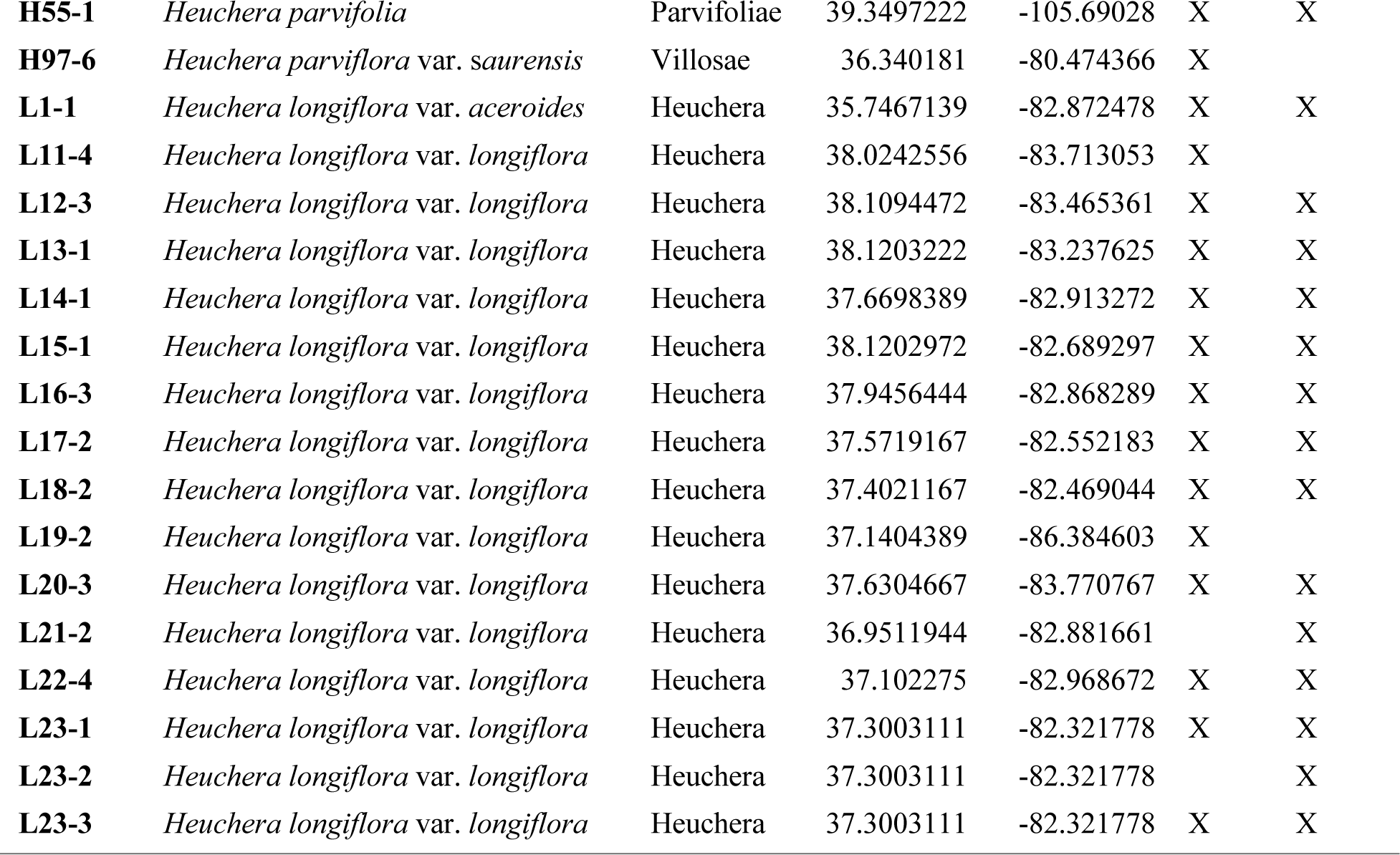
Sampling location and host taxonomy; sample marked “X” according to type of sequencing performed.

**Appendix S2.**
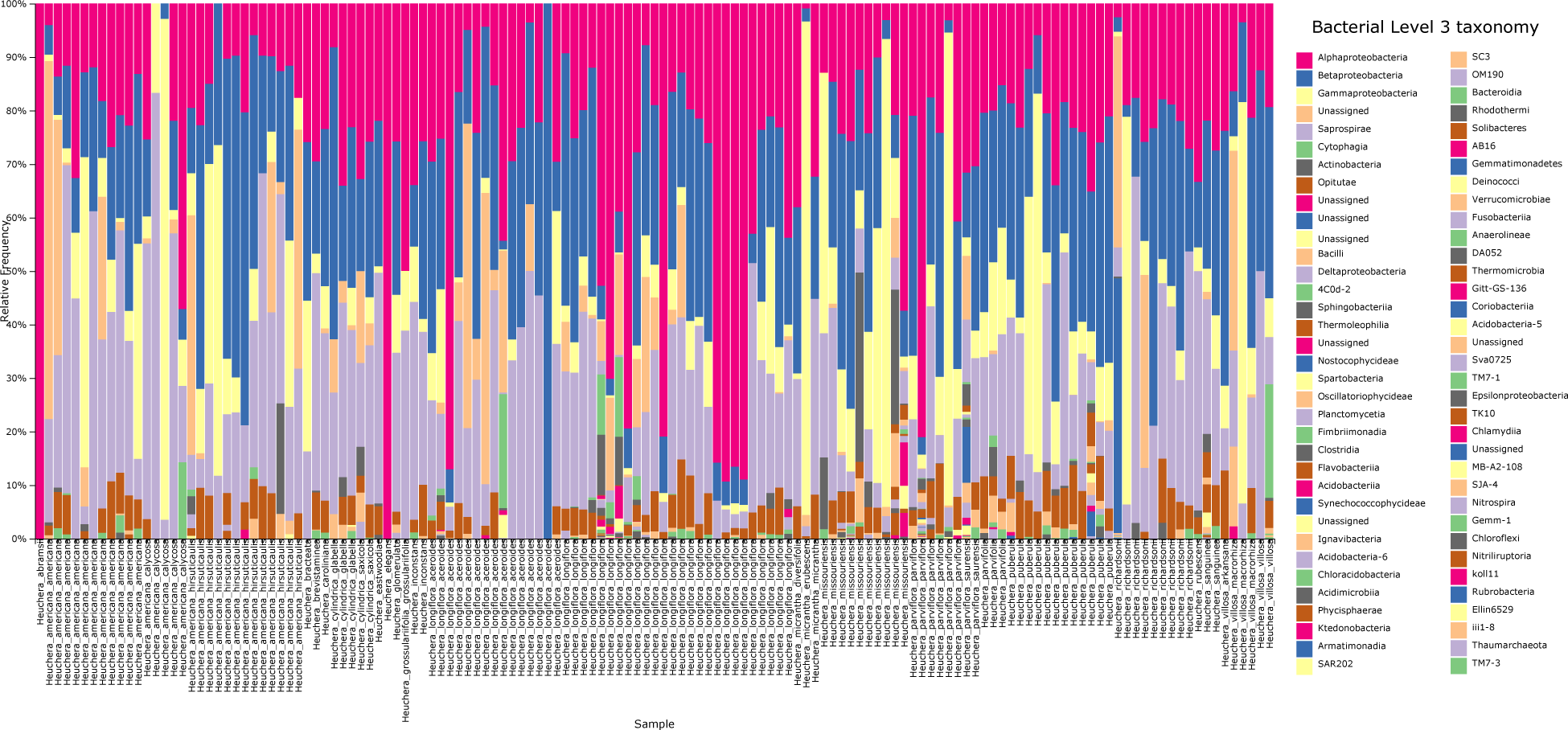
Bacterial class-level diversity and relative abundance per sample.

**Appendix S3.**
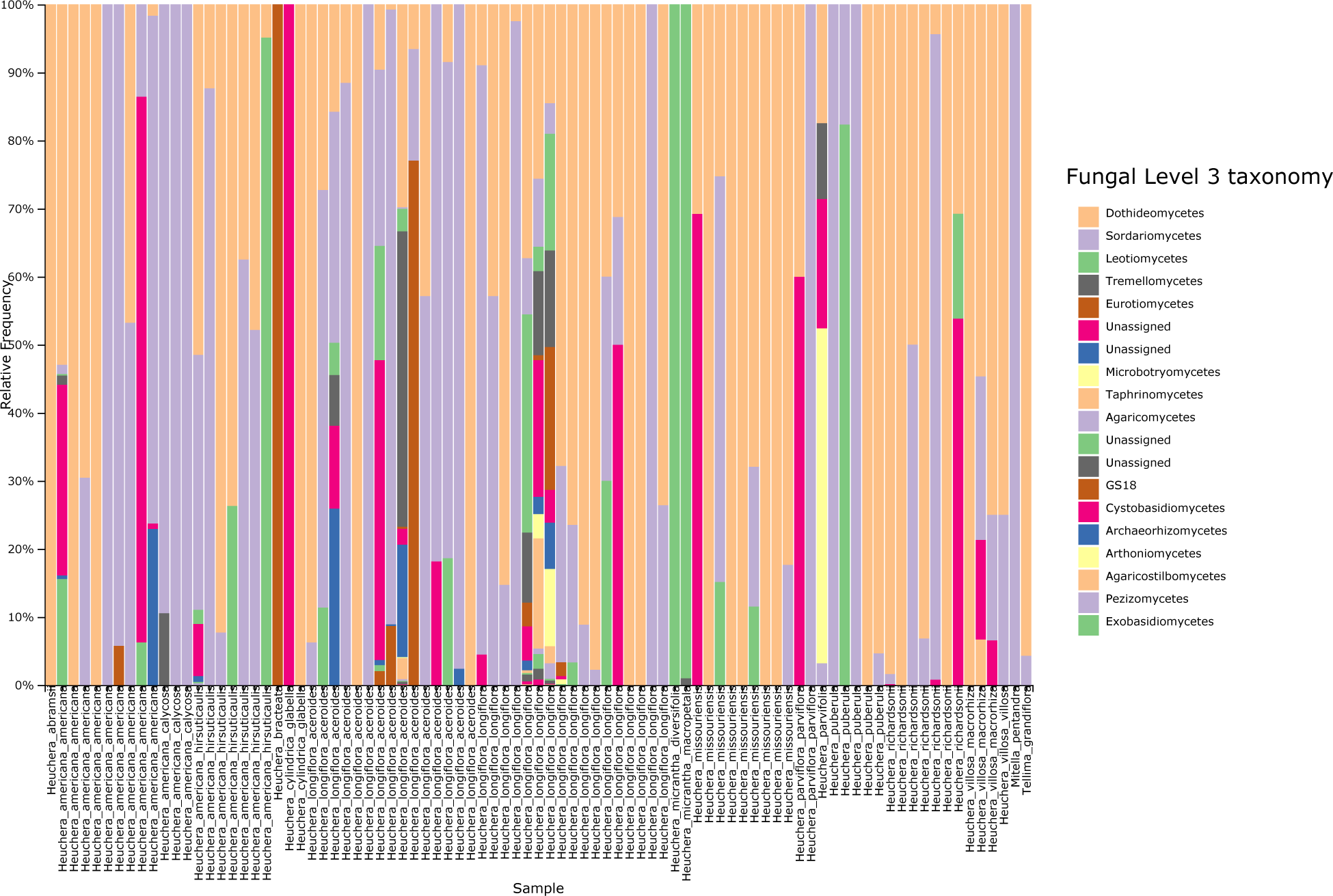
Fungal class-level diversity and relative abundance per sample.

## Literature Cited

Afzal, I., Z. K. Shinwari, S. Sikandar, and S. Shahzad. 2019. Plant beneficial endophytic bacteria: Mechanisms, diversity, host range and genetic determinants. Microbiological Research 221: 36–49.

Afzal, M., S. Yousaf, T. G. Reichenauer, M. Kuffner, and A. Sessitsch. 2011. Soil type affects plant colonization, activity, and catabolic gene expression of inoculated bacterial strains during phytoremediation of diesel. Journal of hazardous Materials 186: 1568–1575.

Andrews, S. 2015. FastQC. Website https://www.bioinformatics.babraham.ac.uk/projects/fastqc/ [accessed 9 October 2019].

Anneberg, T. J., and K. A. Segraves. 2019. Intraspecific polyploidy correlates with colonization by arbuscular mycorrhizal fungi in *Heuchera cylindrica*. American journal of botany 0.

Araújo, W. L., W. Maccheroni, C. I. Aguilar-Vildoso, P. A. Barroso, H. O. Saridakis, and J. L. Azevedo. 2001. Variability and interactions between endophytic bacteria and fungi isolated from leaf tissues of citrus rootstocks. Canadian Journal of Microbiology 47: 229–236.

Arnold, A., and F. Lutzoni. 2007. Diversity and host range of foliar fungal endophytes: are tropical leaves biodiversity hotspots? Ecology 88: 541–549.

Aydogan, E. L., G. Moser, C. Müller, P. Kämpfer, and S. P. Glaeser. 2018. Long-Term Warming Shifts the Composition of Bacterial Communities in the Phyllosphere of Galium album in a Permanent Grassland Field-Experiment. Frontiers in Microbiology 9: 144.

Bahram, M., F. Hildebrand, S. K. Forslund, J. L. Anderson, N. A. Soudzilovskaia, P. M. Bodegom, J. Bengtsson-Palme, et al. 2018. Structure and function of the global topsoil microbiome. Nature 560: 233–237.

Baker, K. L., S. Langenheded, G. W. Nicol, D. Ricketts, K. Killham, C. D. Campbell, and J. I. Prosser. 2009. Environmental and spatial characterisation of bacterial community composition in soil to inform sampling strategies. Soil Biology & Biochemistry 41: 2292– 2298.

Bieker, V. C., F. Sánchez Barreiro, J. A. Rasmussen, M. Brunier, N. Wales, and M. D. Martin. 2020. Metagenomic analysis of historical herbarium specimens reveals a postmortem microbial community. Molecular ecology resources 20: 1206–1219.

Bokati, D., J. Herrera, and R. Poudel. 2016. Soil influences colonization of root-associated fungal endophyte communities of maize, wheat, and their progenitors. Journal of Mycology 2016: 8062073.

Bolyen, E., J. R. Rideout, M. R. Dillon, N. A. Bokulich, C. C. Abnet, G. A. Al-Ghalith, H. Alexander, et al. 2019. Reproducible, interactive, scalable, and extensible microbiome data science using QIIME 2. Nature biotechnology 37: 852–857.

Brigham, L. M., C. P. Bueno de Mesquita, M. J. Spasojevic, E. C. Farrer, D. L. Porazinska, J. G. Smith, S. K. Schmidt, and K. N. Suding. 2023. Drivers of bacterial and fungal root endophyte communities: understanding the relative influence of host plant, environment, and space. FEMS Microbiology Ecology: fiad034.

Bright, M., and S. Bulgheresi. 2010. A complex journey: transmission of microbial symbionts. Nature Reviews Microbiology 8: 218–230.

Bulgarelli, D., K. Schlaeppi, S. Spaepen, E. Ver Loren van Themaat, and P. Schulze-Lefert. 2013. Structure and functions of the bacterial microbiota of plants. Annual review of plant biology 64: 807–838.

Callahan, B. J., P. J. McMurdie, M. J. Rosen, A. W. Han, A. J. A. Johnson, and S. P. Holmes. 2016. DADA2: High-resolution sample inference from Illumina amplicon data. Nature methods 13: 581–583.

Cameron, E. K., I. S. Martins, P. Lavelle, J. Mathieu, L. Tedersoo, M. Bahram, F. Gottschall, et al. 2019. Global mismatches in aboveground and belowground biodiversity. Conservation biology: the journal of the Society for Conservation Biology 33: 1187–1192.

Caporaso, J. G., J. Kuczynski, J. Stombaugh, K. Bittinger, F. D. Bushman, E. K. Costello, N. Fierer, et al. 2010. QIIME allows analysis of high-throughput community sequencing data. Nature methods 7: 335–336.

Christian, N., B. Espino Basurto, A. Toussaint, X. Xu, E. A. Ainsworth, P. E. Busby, and K. D. Heath. 2021. Elevated carbon dioxide reduces a common soybean leaf endophyte. Global Change Biology 27: 4154–4168.

Clay, K. 1990. Fungal Endophytes of Grasses. Annual review of ecology and systematics 21: 275–297.

Coleman-Derr, D., D. Desgarennes, C. Fonseca-Garcia, S. Gross, S. Clingenpeel, T. Woyke, G. North, et al. 2016. Plant compartment and biogeography affect microbiome composition in cultivated and native Agave species. New Phytologist 209: 798–811.

Collado, J., G. Platas, I. González, and F. Peláez. 1999. Geographical and seasonal influences on the distribution of fungal endophytes in *Quercus ilex*. New Phytologist 144: 525–532.

Comby, M., S. Lacoste, F. Baillieul, C. Profizi, and J. Dupont. 2016. Spatial and Temporal Variation of Cultivable Communities of Co-occurring Endophytes and Pathogens in Wheat. Frontiers in microbiology 7: 403.

Copeland, J. K., L. Yuan, M. Layeghifard, P. W. Wang, and D. S. Guttman. 2015. Seasonal community succession of the phyllosphere microbiome. Mol Plant-Microbe Interact 28: 274–285.

Correa-Galeote, D., E. J. Bedmar, and G. J. Arone. 2018. Maize Endophytic Bacterial Diversity as Affected by Soil Cultivation History. Frontiers in microbiology 9: 484.

Daru, B. H., E. A. Bowman, D. H. Pfister, and A. E. Arnold. 2018. A novel proof of concept for capturing the diversity of endophytic fungi preserved in herbarium specimens. Philosophical transactions of the Royal Society of London. Series B, Biological sciences 374.

Dini-Andreote, F. 2020. Endophytes: The Second Layer of Plant Defense. Trends in plant science 25: 319–322.

Ding, T., and U. Melcher. 2016. Influences of Plant Species, Season and Location on Leaf Endophytic Bacterial Communities of Non-Cultivated Plants. PLOS ONE 11: e0150895.

Ding, T., M. W. Palmer, and U. Melcher. 2013. Community terminal restriction fragment length polymorphisms reveal insights into the diversity and dynamics of leaf endophytic bacteria. BMC Microbiology 13.

Douanla-Meli, C., E. Langer, and F. Talontsi Mouafo. 2013. Fungal endophyte diversity and community patterns in healthy and yellowing leaves of *Citrus limon*. Fungal Ecology 6: 212–222.

Eaton, C. J., M. P. Cox, and B. Scott. 2011. What triggers grass endophytes to switch from mutualism to pathogenism? Plant science: an international journal of experimental plant biology 180: 190–195.

Fan, S., L. Miao, H. Li, A. Lin, F. Song, and P. Zhang. 2020. Illumina-based analysis yields new insights into the diversity and composition of endophytic fungi in cultivated Huperzia serrata. PLOS ONE 15: e0242258.

Fang, K., Y.-F. Miao, L. Chen, J. Zhou, Z.-P. Yang, X.-F. Dong, and H.-B. Zhang. 2019. Tissue-Specific and Geographical Variation in Endophytic Fungi of *Ageratina adenophora* and Fungal Associations With the Environment. Frontiers in Microbiology 10: 2919.

Fick, S. E., and R. J. Hijmans. 2017. WorldClim 2: new 1-km spatial resolution climate surfaces for global land areas. International Journal of Climatology 37: 4302–4315.

Fierer, N., and B. Jackson. 2006. The diversity and biogeography of soil bacterial communities. *Proceedings of the National Academy of Sciences*, USA 103: 626–631.

Fisher, P. J., and O. Petrini. 1992. Fungal saprobes and pathogens as endophytes of rice (*Oryza sativa L*.). The New phytologist 120: 137–143.

Fisher, P. J., O. Petrini, and H. M. L. Scott. 1992. The distribution of some fungal and bacterial endophytes in maize (*Zea mays L*.). The New Phytologist 122: 299–305.

Frank, A. C., J. P. Saldierna Guzmán, and J. E. Shay. 2017. Transmission of Bacterial Endophytes. Microorganisms 5: 70.

Folk, R. A. 2015. Biosystematics of the Genus Heuchera (Saxifragaceae). The Ohio State University.

Folk, R. A., and J. V. Freudenstein. 2014. Phylogenetic relationships and character evolution in *Heuchera* (Saxifragaceae) on the basis of multiple nuclear loci. American journal of botany.

Folk, R. A., and J. V. Freudenstein. 2015. ‘Sky islands’ in the eastern USA? — Strong phylogenetic structure in the *Heuchera parviflora* group (Saxifragaceae). Taxon.

Folk, R. A., J. C. Ginori, D. E. Soltis, and A. J. Floden. 2018. Integrative identification of incipient lineages in *Heuchera longiflora* (Saxifragaceae). Botanical journal of the Linnean Society. Linnean Society of London 187: 327–345.

Folk, R. A., J. R. Mandel, and J. V. Freudenstein. 2017. Ancestral gene flow and parallel organellar genome capture result in extreme phylogenomic discord in a lineage of angiosperms. Systematic biology 66: 320–337.

Folk, R. A., C. J. Visger, P. S. Soltis, D. E. Soltis, and R. P. Guralnick. 2018. Geographic range dynamics drove ancient hybridization in a lineage of angiosperms. The American naturalist 192: 171–187.

Gamboa, M. A., and P. Bayman. 2001. Communities of Endophytic Fungi in Leaves of a Tropical Timber Tree (*Guarea guidonia*: Meliaceae). Biotropica 33: 352–360.

Gomes, T., J. A. Pereira, J. Benhadi, T. Lino-Neto, and P. Baptista. 2018. Endophytic and epiphytic phyllosphere fungal communities are shaped by different environmental factors in a mediterranean ecosystem. Microbial Ecology 76: 668–679.

Graham, C. H., D. Storch, and A. Machac. 2018. Phylogenetic scale in ecology and evolution. Global ecology and biogeography: a journal of macroecology 27: 175–187.

Hallmann, J., A. Quadt-Hallmann, W. F. Mahaffee, and J. W. Kloepper. 1997. Bacterial endophytes in agricultural crops. Canadian Journal of Microbiology 43: 895–914.

Hardoim, P. R., L. S. van Overbeek, and J. D. van Elsas. 2008. Properties of bacterial endophytes and their proposed role in plant growth. Trends in microbiology 16: 463–471.

Hardoim, P. R., L. S. van Overbeek, G. Berg, A. M. Pirttilä, S. Compant, A. Campisano, M. Döring, and A. Sessitsch. 2015. The Hidden World within Plants: Ecological and Evolutionary Considerations for Defining Functioning of Microbial Endophytes. Microbiology and Molecular Biology Reviews 79: 293–320.

Harrison, J. G., and E. A. Griffin. 2020. The diversity and distribution of endophytes across biomes, plant phylogeny and host tissues: how far have we come and where do we go from here? Environmental microbiology 22: 2107–2123.

Hengl, T., J. Mendes de Jesus, G. B. M. Heuvelink, M. Ruiperez Gonzalez, M. Kilibarda, A. Blagotić, W. Shangguan, et al. 2017. SoilGrids250m: Global gridded soil information based on machine learning. PloS one 12: e0169748.

Hijmans, R. J., S. E. Cameron, J. L. Parra, P. G. Jones, and A. Jarvis. 2005. Very high resolution interpolated climate surfaces for global land areas. International Journal of Climatology 25: 1965–1978.

Hirano, S. S., L. S. Baker, and C. D. Upper. 1996. Raindrop momentum triggers growth of leaf-associated populations of *Pseudomonas syringae* on field-grown snap bean plants. Appl Environ Microbiol 62: 2560–2566.

Ibarbalz, F. M., N. Henry, M. C. Brandão, S. Martini, G. Busseni, H. Byrne, L. P. Coelho, et al. 2019. Global Trends in Marine Plankton Diversity across Kingdoms of Life. Cell 179: 1084–1097.e21.

Jin, H., Z. Yan, Q. Liu, X. Yang, J. Chen, and B. Qin. 2013. Diversity and dynamics of fungal endophytes in leaves, stems and roots of *Stellera chamaejasme* L. in northwestern China. Antonie Van Leeuwenhoek 104: 949–963.

Johnson, G. 2019. High throughput DNA extraction of legume root nodules for rhizobial metagenomics. AMB Express 9: 47.

Jumpponen, A., and J. M. Trappe. 1998. Dark septate endophytes: a review of facultative biotrophic root-colonizing fungi. The New phytologist 140: 295–310.

Jung, J. H., F. Reis, C. L. Richards, and O. Bossdorf. 2021. Understanding plant microbiomes requires a genotype × environment framework. American journal of botany 108: 1820– 1823.

Kandel, S. L., P. M. Joubert, and S. L. Doty. 2017. Bacterial Endophyte Colonization and Distribution within Plants. Microorganisms 5: 77.

Khare, E., J. Mishra, and N. K. Arora. 2018. Multifaceted Interactions Between Endophytes and Plant: Developments and Prospects. Frontiers in microbiology 9: 2732.

Koide, R., K. D. Ricks, and E. R. Davis. 2017. Climate and dispersal influence the structure of leaf fungal endophyte communities of Quercus gambelii in the eastern Great Basin, USA. Fungal Ecology 30: 19–28.

Kozich, J. J., S. L. Westcott, N. T. Baxter, S. K. Highlander, and P. D. Schloss. 2013. Development of a dual-index sequencing strategy and curation pipeline for analyzing amplicon sequence data on the MiSeq Illumina sequencing platform. Applied and environmental microbiology 79: 5112–5120.

Langenfeld, A., S. Prado, B. Nay, C. Cruaud, S. Lacoste, E. Bury, F. Hachette, et al. 2013. Geographic locality greatly influences fungal endophyte communities in *Cephalotaxus harringtonia*. Fungal Biology 117: 124–136.

Larran, S., A. Perelló, M. R. Simón, and V. Moreno. 2002. Isolation and analysis of endophytic microorganisms in wheat (*Triticum aestivum* L.) leaves. World journal of microbiology & biotechnology 18: 683–686.

Lau, M. K., A. E. Arnold, and N. C. Johnson. 2013. Factors influencing communities of foliar fungal endophytes in riparian woody plants. Fungal Ecology 6: 365–378.

Leuchtmann, A. 1992. Systematics, distribution, and host specificity of grass endophytes. Natural toxins 1: 150–162.

Matsumura, E., and K. Fukuda. 2013. A comparison of fungal endophytic community diversity in tree leaves of rural and urban temperate forests of Kanto district, eastern Japan. Fungal Biology 117: 191–201.

McDonald, D., M. N. Price, J. Goodrich, E. P. Nawrocki, T. Z. DeSantis, A. Probst, G. L. Andersen, et al. 2012. An improved Greengenes taxonomy with explicit ranks for ecological and evolutionary analyses of bacteria and archaea. The ISME journal 6: 610–618.

Middleton, N. D. S. G. Thomas, and United Nations Environment Programme. 1992. World atlas of desertification.

Miliute, I., O. Buzaite, D. Baniulis, and V. Stanys. 2015. Bacterial endophytes in agricultural crops and their role in stress tolerance: a review. Zemdirbyste-Agriculture 102: 465–478.

Mina, D., J. A. Pereira, T. Lino-Neto, and P. Baptista. 2020. Epiphytic and Endophytic Bacteria on Olive Tree Phyllosphere: Exploring Tissue and Cultivar Effect. Microbial Ecology 80: 145–157.

Nilsson, R. H., K.-H. Larsson, A. F. S. Taylor, J. Bengtsson-Palme, T. S. Jeppesen, D. Schigel, P. Kennedy, et al. 2019. The UNITE database for molecular identification of fungi: handling dark taxa and parallel taxonomic classifications. Nucleic acids research 47: D259–D264.

O’Brien, A. M., N. A. Ginnan, M. Rebolleda-Gómez, and M. R. Wagner. 2021. Microbial effects on plant phenology and fitness. American journal of botany 108: 1824–1837.

Pang, B., D. Yin, Y. Zhai, A. He, L. Qiu, Q. Liu, N. Ma, et al. 2022. Diversity of endophytic fungal community in *Huperzia serrata* from different ecological areas and their correlation with Hup A content. BMC Microbiology 22: 191.

Pedregosa, F., G. Varoquaux, A. Gramfort, V. Michel, B. Thirion, O. Grisel, M. Blondel, et al. 2011. Scikit-learn: Machine Learning in Python. Journal of machine learning research: JMLR 12: 2825–2830.

Penner, S., and Y. Sapir. 2021. Foliar endophytic fungi inhabiting an annual grass along an aridity gradient. Current Microbiology 78: 2080–2090.

QGIS Development Team. 2021. QGIS Geographic Information System. Open Source Geospatial Foundation. URL https://www.qgis.org

R Core Team. 2021. R: A language and environment for statistical computing. R Foundation for Statistical Computing, Vienna, Austria. URL https://www.R-project.org/.

Rodriguez, R. J., J. F. White Jr, A. E. Arnold, and R. S. Redman. 2009. Fungal endophytes: diversity and functional roles. New Phytologist 182: 314–330.

Romero, A., G. Carrion, and V. Rico-Gray. 2001. Fungal latent pathogens and endophytes from leaves of. Fungal Diversity 7: 81–87.

Rosenblueth, M., and E. Martínez-Romero. 2006. Bacterial Endophytes and Their Interactions with Hosts. Molecular Plant-Microbe Interactions® 19: 827–837.

Schardl, C. L. 2001. *Epichloë festucae* and related mutualistic symbionts of grasses. Fungal genetics and biology: FG & B 33: 69–82.

Schardl, C. L., and H. F. Tsai. 1992. Molecular biology and evolution of the grass endophytes.Natural toxins 1: 171–184.

Schmidt, P.-A., M. Bálint, B. Greshake, C. Bandow, J. Römbke, and I. Schmitt. 2013. Illumina metabarcoding of a soil fungal community. Soil biology & biochemistry 65: 128–132.

Shakya, M., N. Gottel, H. Castro, Z. K. Yang, L. Gunter, J. Labbé, W. Muchero, et al. 2013. A multifactor analysis of fungal and bacterial community structure in the root microbiome of mature Populus deltoides trees. PloS One 8: e76382.

Shen, C., A. Gunina, Y. Luo, J. Wang, J.-Z. He, Y. Kuzyakov, A. Hemp, et al. 2020. Contrasting patterns and drivers of soil bacterial and fungal diversity across a mountain gradient. Environmental Microbiology 22: 3287–3301.

de Souza, R. S. C., V. K. Okura, J. S. L. Armanhi, B. Jorrín, N. Lozano, M. J. da Silva, M. González-Guerrero, et al. 2016. Unlocking the bacterial and fungal communities assemblages of sugarcane microbiome. Scientific Reports 6: 28774.

Stone, B. W. G., and C. R. Jackson. 2019. Canopy position is a stronger determinant of bacterial community composition and diversity than environmental disturbance in the phyllosphere. FEMS Microbiol Ecol 95:1–11. 95: 1–11.

Stone, B. W. G., and C. R. Jackson. 2021. Seasonal patterns contribute more towards phyllosphere bacterial community structure than short-term perturbations. Microbial Ecology 81: 146–156.

Tellez, P. H., A. E. Arnold, A. B. Leo, K. Kitajima, and S. A. Van Bael. 2022. Traits along the leaf economics spectrum are associated with communities of foliar endophytic symbionts. Frontiers in Microbiology 13.

Thompson, L. R., J. G. Sanders, D. McDonald, A. Amir, J. Ladau, K. J. Locey, R. J. Prill, et al. 2017. A communal catalogue reveals Earth’s multiscale microbial diversity. Nature 551: 457–463.

Title, P. O., and J. B. Bemmels. 2018. ENVIREM: an expanded set of bioclimatic and topographic variables increases flexibility and improves performance of ecological niche modeling. Ecography 41: 291–307.

Trivedi, P., J. E. Leach, S. G. Tringe, T. Sa, and B. K. Singh. 2020. Plant–microbiome interactions: from community assembly to plant health. Nature reviews. Microbiology.

Van Bael, S., C. Estrada, and A. E. Arnold. 2017. Foliar endophyte communities and leaf traits in tropical trees. The fungal community: its organization and role in the ecosystem, 79–94. CRC Press, Boca Raton, FL, USA.

Vorholt, J. A. 2012. Microbial life in the phyllosphere. Nature Reviews Microbiology 10: 828– 840.

Wei, Y., G. Lan, Z. Wu, B. Chen, F. Quan, M. Li, S. Sun, and H. Du. 2022. Phyllosphere fungal communities of rubber trees exhibited biogeographical patterns, but not bacteria. Environmental Microbiology 24: 3769–3782.

Wells, E. F. 1984. A Revision of the Genus *Heuchera* (Saxifragaceae) in Eastern North America. Systematic Botany Monographs 3: 45.

Wemheuer, F., D. Berkelmann, B. Wemheuer, R. Daniel, S. Vidal, and H. B. Bisseleua Daghela. 2020. Agroforestry Management Systems Drive the Composition, Diversity, and Function of Fungal and Bacterial Endophyte Communities in *Theobroma cacao* Leaves. Microorganisms 8: 405.

Wemheuer, F., B. Wemheuer, R. Daniel, and S. Vidal. 2019. Deciphering bacterial and fungal endophyte communities in leaves of two maple trees with green islands. Scientific Reports 9: 14183.

Yang, T., L. Tedersoo, P. S. Soltis, D. E. Soltis, J. A. Gilbert, M. Sun, Y. Shi, et al. 2019. Phylogenetic imprint of woody plants on the soil mycobiome in natural mountain forests of eastern China. The ISME journal 13: 686–697.

Yang, X., P. Wang, B. Xiao, Q. Xu, Q. Guo, S. Li, L. Guo, et al. 2023. Different assembly mechanisms of leaf epiphytic and endophytic bacterial communities underlie their higher diversity in more diverse forests. Journal of Ecology 00: 1–12.

Yeoh, Y. K., P. G. Dennis, C. Paungfoo-Lonhienne, L. Weber, R. Brackin, M. A. Ragan, S. Schmidt, and P. Hugenholtz. 2017. Evolutionary conservation of a core root microbiome across plant phyla along a tropical soil chronosequence. Nature communications 8: 215.

Zarraonaindia, I., S. M. Owens, P. Weisenhorn, K. West, J. Hampton-Marcell, S. Lax, N. A. Bokulich, et al. 2015. The Soil Microbiome Influences Grapevine-Associated Microbiota J. K. Jansson [ed.],. mBio 6: e02527–14.

Zhang, G., G. Wei, F. Wei, Z. Chen, M. He, S. Jiao, Y. Wang, et al. 2021. Dispersal limitation plays stronger role in the community assembly of fungi relative to bacteria in rhizosphere across the arable area of medicinal plant. Front Microbiol 12: 713523.

Zhang, T., and Y.-F. Yao. 2015. Endophytic Fungal Communities Associated with Vascular Plants in the High Arctic Zone Are Highly Diverse and Host-Plant Specific. PLOS ONE 10: e0130051.

Zhou, Y., Y. Wei, M. Ryder, H. Li, Z. Zhao, R. Toh, P. Yang, et al. 2023. Soil salinity determines the assembly of endophytic bacterial communities in the roots but not leaves of halophytes in a river delta ecosystem. Geoderma 433: 116447.

Zimmerman, N. B., and P. M. Vitousek. 2012. Fungal endophyte communities reflect environmental structuring across a Hawaiian landscape. PNAS 109: 13022–13027.

